# Epigenetic potential and dispersal propensity in a free-living songbird: a spatial and temporal approach

**DOI:** 10.1101/2024.11.07.622405

**Authors:** Blanca Jimeno, Marianthi Tangili, Julio C. Domínguez, David Canal, Carlos Camacho, Jaime Potti, Jesús T. García, Jesús Martínez-Padilla, Mark Ravinet

## Abstract

Natal dispersal is a key life-history trait determining fitness and driving population dynamics, genetic structure and species distributions. Despite existing evidence that not all phenotypes are equally likely to successfully establish in new areas, the mechanistic underpinnings of natal dispersal remain poorly understood. The propensity to disperse into a new environment can be favored by a high degree of phenotypic plasticity which facilitates local adaptation and may be achieved via epigenetic mechanisms, which modify gene expression and enable rapid phenotypic changes. Epigenetic processes occur in particular genomic regions - DNA methylation on CpG sites in vertebrates -, and thus individual genomes may differ in their capacity to be modified epigenetically. This “Epigenetic potential” (EP) may represent the range of phenotypic plasticity attainable by an individual, and be a key determinant of successful settlement in novel areas. We investigated the association between EP – quantified as the number of genome-wide CpG variants – and natal dispersal propensity in a long-term study population of Pied flycatchers (*Ficedula hypoleuca*) monitored since colonization of a new habitat 35 years ago. We tested this association at three levels, comparing EP between: i) individuals dispersing between and within habitat patches; ii) immigrants to the population and locally-born individuals; and iii) individuals from first (comprising colonizers or their direct descendants) and later generations of the population (consisting of locally-born individuals, which did not show natal dispersal). Results show a significant, positive association between EP and dispersal propensity in comparisons i) and iii), but not ii). Furthermore, CpG variants were non-randomly distributed across the genome, suggesting species- and/or population-specific CpGs being more frequent in promoters and exons. Our findings point to EP playing a role in dispersal propensity at spatial and temporal scales, supporting the idea that epigenetically-driven phenotypic plasticity facilitates dispersal and environmental coping in free-living birds.

## 1. INTRODUCTION

In an increasingly changing world, organisms need to adjust to environmental fluctuations, and life history traits should evolve towards maximizing fitness in the face of environmental variability and physiological trade-offs. In changing environments, the evolution of life history traits by natural selection will depend on genetically-determined phenotypic diversity, which provides the variation needed for selection to drive adaptive changes (Martínez-Padilla *et al*. 2017; Siepielski *et al*. 2017; Teplitsky *et al*. 2014; Bell 2010). Furthermore, in a time when environments are changing so rapidly, individuals are forced to adjust to prevailing conditions, and can respond quickly through phenotypic plasticity, which will depend on the environmental conditions and does not require genetic change (Pigliucci 2005). Understanding the mechanisms driving organismal responses to environmental change and the evolutionary relevance of phenotypic plasticity becomes fundamental in predicting the biological impact of ongoing climate and land-use changes. Epigenetic mechanisms are changes in gene expression that do not involve changes in the DNA sequence and potentially enable rapid and adaptive responses via increased phenotypic plasticity. Understanding the role of these mechanisms in local adaptation of wild populations under current environmental changes is becoming increasingly important.

Natal dispersal - the movement between the place of birth and first breeding - is a particularly relevant life history trait affecting most ecological and evolutionary processes determining individual fitness (Clobert *et al*. 2009), and potentially scaling up into population dynamics, genetic structure and species distributions (Johnson *et al*. 1990; Whitlock 2001). Natal dispersal comprises departure (initiation to leave the natal patch), transfer (movement), and settlement (establishment in the breeding patch), and is often influenced by multiple morphological, physiological and behavioural traits (Clobert *et al*. 2009). Previous research on natal dispersal shows that dispersers are not a random sample of the source population (Bowler & Benton 2005; Clobert *et al*. 2009; Edelaar & Bolnick 2012; Camacho *et al*. 2013), and may share traits that facilitate settlement, mitigating the costs of dispersal (Camacho *et al*. 2013; Barbraud *et al*. 2003; Duckworth 2008). However, despite existing evidence that not all phenotypes are equally likely to disperse and establish in new areas (Bowler & Benton 2005; Clobert *et al*. 2009; Edelaar & Bolnick 2012), the mechanistic underpinnings of natal dispersal propensity remain poorly understood, partly due to the scarcity of longitudinal and experimental data (Dingemanse *et al*. 2003). Studies so far point at variation in natal dispersal resulting from the interplay between external factors (i.e., environmental cues (Brown & Brown 1992; Delgado *et al*. 2010) and individual phenotype (e.g., body size, coloration or personality (Dingemanse *et al*. 2003; Barbraud *et al*. 2003; Delgado *et al*. 2010; Camacho *et al*. 2013; Camacho *et al*. 2017b). Dispersal phenotypes has a detectable genetic basis - along with a complex genetic architecture involving many genes - in many organisms, including birds (reviewed by Saastamoinen *et al*. 2018, Hansson *et al*. 2003; Doligez *et al*. 2009; Korsten *et al*. 2013). Heritability studies suggest that much of the phenotypic variation linked to dispersal is explained by environmental variation, which entails that the adaptive potential of dispersal may be context-dependent (Saatoglu *et al*. 2024, but see Charmantier *et al*. 2011). Furthermore, there is also evidence demonstrating strong epigenetic signals in dispersal propensity (van Petegem *et al*., 2015).

Nevertheless, insights of the genetic and environmental factors underlying specific traits affecting individual variation in natal dispersal behaviour are scarce. Thus, investigating which mechanisms underlie variation in dispersal phenotypes and the associated interaction between genes and the environment is crucial to predicting whether populations harbor adaptive potential for dispersal, as well as understanding the eco-evolutionary dynamics of this important trait (Saatoglu *et al*. 2024).

Optimizing dispersal decisions according to environmental factors and individual phenotype usually requires a high degree of phenotypic plasticity that allows individuals to respond adaptively when facing novel environments (Duckworth 2008; Kilvitis *et al*. 2017; but see Edelaar *et al*. 2017; Fig. 1). Epigenetic mechanisms link environmental variation to phenotypic plasticity by modifying gene expression (Smith & Meissner 2013), enabling more rapid phenotypic changes than are possible via genetic adaptation. These mechanisms may allow genotypes to mask and/or manifest phenotypic plasticity reversibly within generations, acting as a ‘buffer’ in the face of rapid environmental change. The evolutionary impact of epigenetic mechanisms, however, remains unclear, partly due to the uncertainty over how epigenetic marks are inherited through the germ line in animals (Sepers *et al*. 2019). Epigenetic variation predominates in particular genomic regions, and thus individual genomes may differ in their capacity to be modified epigenetically (Kilvitis *et al*. 2017; Smith & Meissner 2013; Feinberg & Irizarry 2010), namely “Epigenetic Potential” (i.e. susceptibility of a genome (or genome sequence, especially promoters) to show epigenetic changes; Kilvitis *et al*. 2017; Hanson *et al*. 2020, 2021, 2022). One of the most studied epigenetic processes is DNA methylation. It consists of the addition of a methyl group to the DNA sequence, and has been suggested as a pivotal source of phenotypic plasticity in several animal species (Feinberg & Irizarry 2010; Angers *et al*. 2010; Sepers *et al*. 2019; Sammarco *et al*. 2022). DNA methylation in vertebrates occurs predominantly in cytosines that occur before guanines (i.e. CpG sites). Hence, a higher number of CpG sites in the genome may be translated in a higher susceptibility to show DNA methylation / demethylation and thus epigenetically-driven phenotypic plasticity (Feinberg & Irizarry 2010; Kilvitis *et al*. 2017; Hanson *et al*. 2022). Moreover, if epigenetic potential represents the range of phenotypic plasticity achievable to an individual, it may be beneficial for dispersing individuals when coping with novel environments (Kilvitis *et al*. 2017; Hanson *et al*. 2020; Hanson *et al*. 2022). Furthermore, because epigenetic potential should be a genetic, heritable trait, it may represent a link between genetic inheritance and epigenetic processes, enhancing its evolutionary impact. In line with these predictions, recent evidence suggests that a high epigenetic potential (quantified as CpG abundance) may enable house sparrow (*Passer domesticus*) populations to thrive throughout environmental variability and colonize new areas. Introduced and expanding sparrow populations exhibited higher epigenetic potential in specific genome regions (Hanson *et al*. 2022), including two immune gene promoters (Hanson *et al*. 2020), when compared with historic ones, and results suggested epigenetic potential being favored by selection in recently colonized areas (Hanson *et al*. 2022). On the basis of this evidence, the level of epigenetically-driven phenotypic plasticity may underlie variation in (natal) dispersal, such that individuals showing higher dispersal propensity and colonization success will also show higher epigenetic potential.

**Figure 1.**
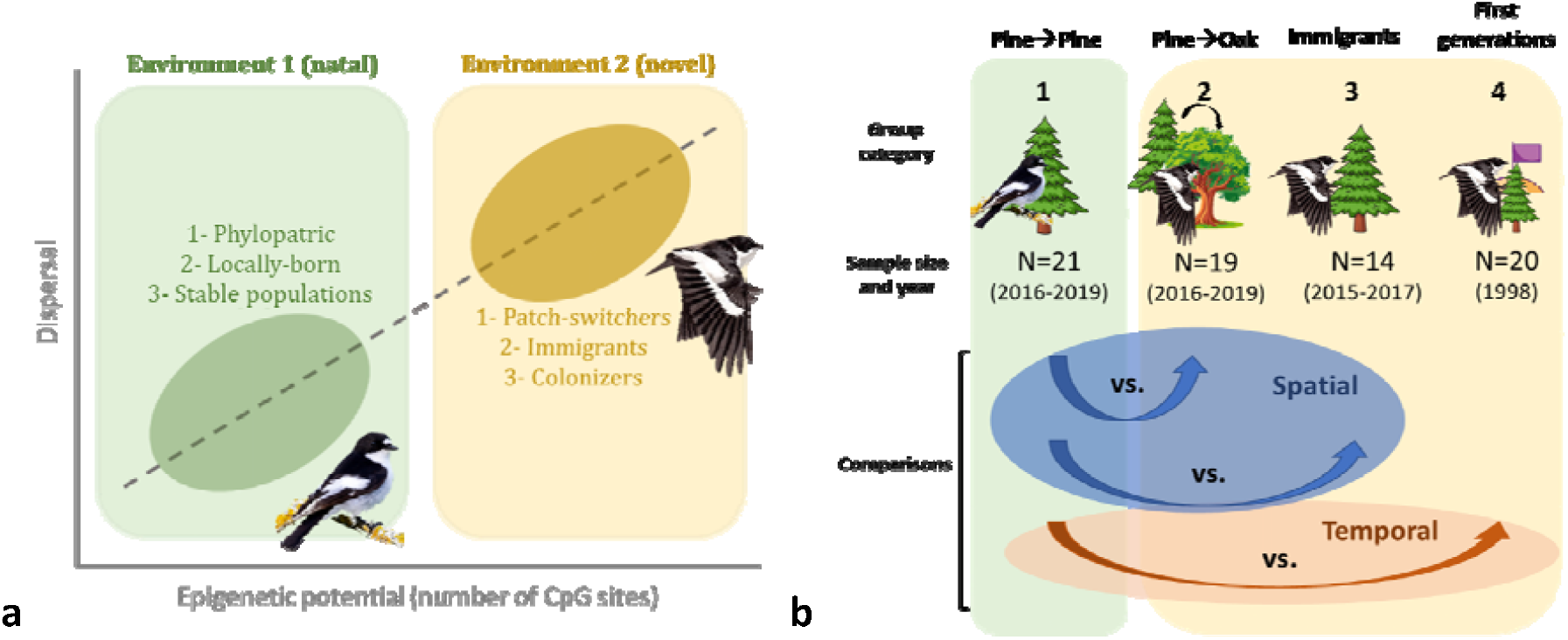
(a) Schematic overview of the predicted positive association between epigenetic potential (as a source of phenotypic plasticity) and natal dispersal. Note that dispersal categories linked to high (orange) or low (green) epigenetic potential relate to each of the three comparisons included in the study, and correspond to comparisons at different spatial (1,2) and temporal (3) scales. (b) Schematic representation of the group categories and comparisons included in the study, along with sample size and sampling years. Different group categories correspond to: 1. Birds reared and breeding in the pine forest, 2. Birds reared in the pine forest but breeding in the oak forest, 3. Birds identified as immigrants to the population, 4. First generations of birds breeding in the pine forest after patch colonization. For further details see methods and Table 1.

Because epigenetic potential research in ecology is in its infancy, there are still many unanswered questions regarding the interpretation of this trait and its evolutionary relevance. Importantly, the functional mechanisms linking CpG abundance and regulation of gene expression are still under debate, mostly because the molecular dynamics of CpG sites and CpG islands do not always support a direct association between CpG number and the susceptibility for environmentally-driven regulation of gene expression (Chen *et al*. 2018; Lelli *et al*. 2012). Furthermore, previous studies quantifying epigenetic potential have targeted specific genes and regions (i.e. promoters) (e.g. Hanson *et al*. 2020, 2022), and therefore a key next step is to test whether the above associations hold when looking at the whole genome. This step becomes critical in the context of dispersal behavior, a complex trait in which many genes are likely involved, and genetic and environmental factors interact (Saastamoinen *et al*. 2018, Saatoglu *et al*. 2024). Investigating the role of epigenetic potential in ecologically relevant traits - such as natal dispersal - in the face of environmental change is crucial in understanding its eco-evolutionary consequences.

**Table 1:**
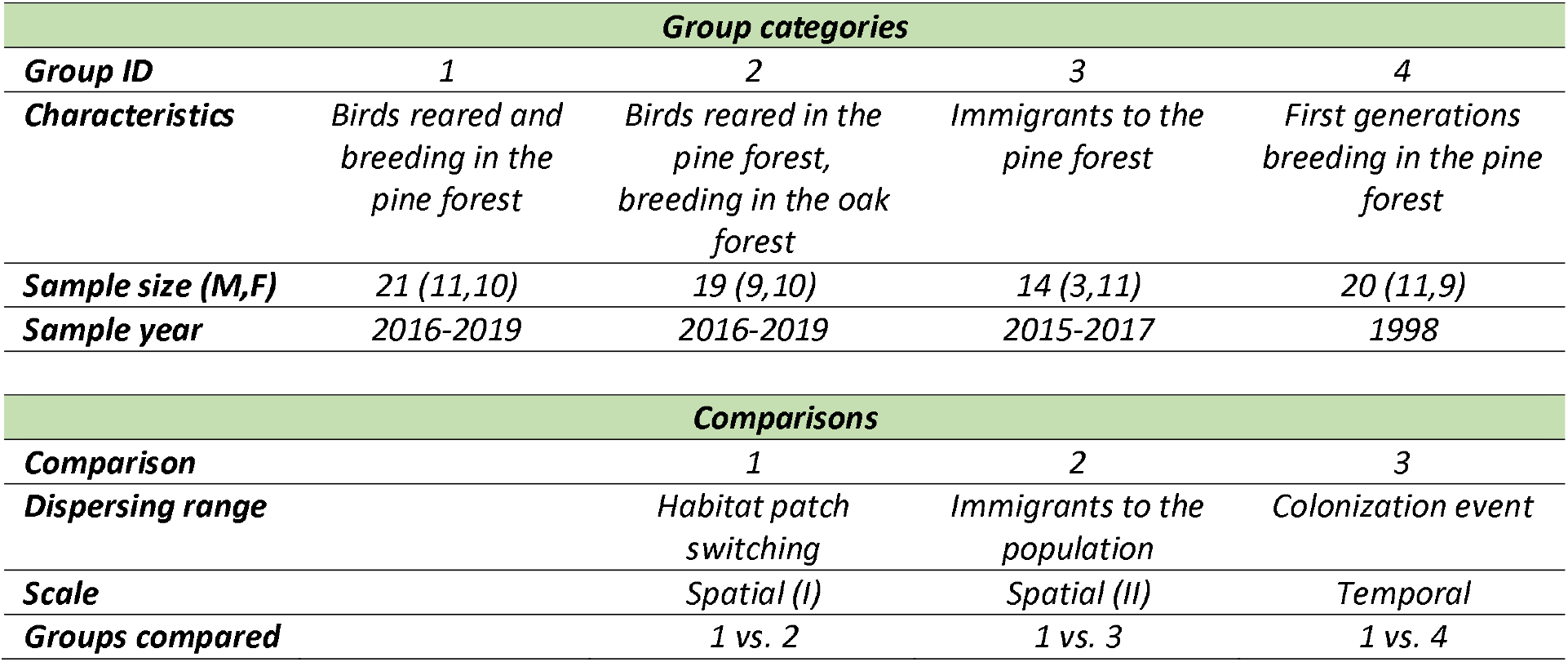
Group categories and comparisons included in the study. Group categories are shown along with sample sizes (including males and females) and years. Comparisons also appear grouped by the scale (i.e. spatial or temporal).

| **Group categories** |  |  |  |  |
| --- | --- | --- | --- | --- |
| **Group ID** | **1** | **2** | **3** | **4** |
| **Characteristics** | *Birds reared and breeding in the pine forest* | *Birds reared in the pine forest, breeding in the oak forest* | *Immigrants to the pine forest* | *First generations breeding in the pine forest* |
| **Sample size (M,F)** | 21 (11,10) | 19 (9,10) | 14 (3,11) | 20 (11,9) |
| **Sample year** | 2016-2019 | 2016-2019 | 2015-2017 | 1998 |

| **Comparisons** |  |  |  |
| --- | --- | --- | --- |
| **Comparison** | **1** | **2** | **3** |
| **Dispersing range** | *Habitat patch switching* | *Immigrants to the population* | *Colonization event* |
| **Scale** | *Spatial (I)* | *Spatial (II)* | *Temporal* |
| **Groups compared** | *1 vs. 2* | *1 vs. 3* | *1 vs. 4* |

In this work, we tested the prediction that epigenetic potential is associated with natal dispersal propensity, by scaling it up from individual life history decisions to a colonization event using a long-term study population of wild Pied flycatchers (*Ficedula hypoleuca*). Natal dispersal of birds originating from this population has been monitored for four decades and, crucially, since the first settlers arrived in one of the population patches (Camacho *et al*. 2019). We performed three different comparisons using historical and newly collected samples from individuals differing in dispersal behavior (Fig. 1b) to investigate the association between epigenetic potential, quantified as number of CpG variants, and dispersal patterns at both spatial and temporal scales. Our general prediction was that more dispersive groups will show higher epigenetic potential (Fig. 1a). We further annotated CpG variants, hypothesizing that their distribution in functional categories across the genome, if involved in environmental coping and adaptation, will not be random.

## 2. METHODS

### 2.1. Study system and population monitoring

Samples were collected in population of pied flycatchers breeding in nest boxes in central Spain (La Hiruela, 41°04’ N, 3°27’ W; 1250 masl), and monitored since 1984. Pied flycatchers are small passerines and long-distance migrants breeding across Eurasia. Individuals show limited natal dispersal (median dispersal distance, males: 445 m, females: 600 m; Potti & Montalvo 1991). Breeding dispersal (i.e. dispersal between breeding sites) was significantly more restricted (ca. 2%) than natal dispersal (ca. 27% between breeding patches), especially in older breeders, which mostly return to bred at the same site as in previous years or its immediate vicinity (Harvey *et al*. 1984; Montalvo & Potti 1992).

The study population includes two plots located in different habitats, a deciduous oak forest (*Quercus pyrenaica*) of 9.3 ha and a mixed coniferous plantation (mostly *Pinus sylvestris*) of 4.8 ha, 1.1 km apart. Before the onset of the long-term study, a small population of flycatchers was already present in the oak forest (0.13 pairs/Ha), and numbers increased after nest boxes were added in 1984 (up to 12.7 pairs/Ha; Camacho *et al*. 2013; Potti & Montalvo 1990; Morales-Mata *et al*. 2022). In contrast, Pied flycatchers did not breed in the pine plantation before the study began due to the lack of natural cavities, and they only established after nest boxes were provided in 1988. Bird numbers in the pine forest increased steeply since the first pair established in 1988, but it was not until 1993 that breeding density became comparable to that in the oak forest. Based on this, we defined the period between 1988 and 1993 as the colonization phase (Camacho *et al*. 2019. Since 1995, the number and location of nest boxes remain unchanged, with 156 nest boxes in the oak forest and 83 nest boxes in the pine forest.

Every year, during the breeding season from around the third week of April (when first males arrive from migration) to the beginning of July, all breeding adults and offspring are captured, identified with metal rings, and measured for standard morphological traits (e.g. body size, weight, sexual traits); life history traits, such as lying date, clutch size or fledgling number, are also assessed. Adults were captured with a nest-box trap (Friedman *et al*., 2008) while feeding nestlings on day 8-10 posthatching. Chicks were measured, ringed and blood-sampled on day 13 post-hatching. Since year 2005, all captured individuals are blood sampled, and samples are stored in ethanol at −20 °C until analysis. The study area and field protocols have been described in detail in Camacho 2018; Potti *et al*., 2018. The current protocols maximize the capacity to identify a high proportion of breeding individuals (Camacho *et al*. 2017a). Local recruitment rates are among the highest reported on this species (∼13%; Camacho *et al*. 2016), and so natal dispersal of most breeding adults is known. ∼170 breeding pairs are monitored every year and >19000 nestlings have been banded in the population so far, of which ∼2000 became breeders (recruits). Unringed birds first caught as breeding adults were considered immigrants, whereas local recruits ringed as nestlings are hereafter referred to as ‘locally-born’. The average proportion of individuals identified as immigrants (i.e. coming from outside the study population) is 53.3% (47.9%% males, 58.0% females – unpublished data). Unringed (immigrant) individuals were aged as first year or older based on their plumage traits (Potti & Montalvo 1991).

### 2.2. Sample selection and group categories

We focused on the pine patch for sample selection, as it has been monitored since the establishment of the population. We selected blood samples for sequencing from the historical database to test for the association between natal dispersal propensity and epigenetic potential at 3 different scales (two spatial, one temporal), taking the pine patch as reference (Table 1,Fig. 1b):

- Habitat patch switching (Spatial I): We compared epigenetic potential of individuals that originated from the pine forest and returned there to breed (Group 1), and that of individuals from the pine forest, but that dispersed to the oak forest to breed for the first time (Group 2).
- Immigrants to the population (Spatial II): We compared epigenetic potential of individuals that originated from the pine forest and returned there to breed (Group 1), and that of individuals categorized as immigrants, which inherently dispersed from the place of birth (Group 3).
- Colonization event (Temporal): We compared epigenetic potential of individuals that originated from the pine forest and returned there to breed in the last period of the study (Group 1), and that of individuals breeding in the pine forest right after the colonization event (Group 4). These latter individuals were either first colonizers of the pine forest or their direct descendants. Thus, they either dispersed to the pine forest through natal dispersal, or inherited epigenetic potential (CpG sites) from their parents who did.

We performed sample selection so that samples from groups 1 (faithful to the pine patch), 2 (patch switchers) and 3 (immigrants to the pine patch), representing the current population, covered a relatively short time period (< 5 years) to minimize temporal effects. Samples from group 4 (individuals from first generations of the population) date from 1998, the oldest available samples for the study population. Sample selection also guaranteed that individuals were not family related (i.e. parents and their offspring, or brothers / sisters). We aimed for 20 individuals per group, but due to poor DNA quality and / or sequencing failure, our final sample sizes were a bit lower for some of the groups (see Table 1,Fig. 1b).

### 2.3. DNA extraction and sequencing

DNA from adult breeders (recruits and immigrants) in recent years had already been obtained as part of other ongoing projects. Total genomic DNA was extracted from blood samples using a standard ammonium acetate protocol (Strauss, 1998), and diluted to a working concentration of 25 ng/μL. Besides, on an additional set of samples from 1998 (used in one of the group comparisons; see below), DNA had already been extracted from whole blood using Chelex resin-based extraction (Walsh *et al*. 1991; Potti *et al*. 2002).

Whole genome sequencing was performed by Novogene Europe in two sequencing runs, where paired-end Illumina next-generation sequencing (150bp) was carried out on an Illumina NovaSeq 6000 sequencer. Libraries were prepared using either the Novogene NGS DNA Library Prep Set (Cat No.PT004) at Novogene, or small-scale library prep (1/10th standard volumes) for Illumina DNA Prep with IDT index sets, using the Mosquito aliquoting system at the DeepSeq Facility at the University of Nottingham, UK. Both library preparation methods have previously been validated and shown to have comparable sequencing outcomes in another passerine species, the house sparrow (M. Ravinet, unpubl. data).

### 2.4. Bioinformatic processing

Genome sequences were aligned against the collared flycatcher (Ficedula albicollis) reference genome (https://www.ncbi.nlm.nih.gov/datasets/genome/GCF_000247815.1/; accession AGTO00000000.2), the closest relative species with a reference genome available. In order to map reads to the collared flycatcher reference genome, we used a custom genotyping pipeline developed in Nextflow (https://github.com/markravinet/genotyping_pipeline). The pipeline is based on a previously used protocol (Ravinet *et al*. 2018) and is designed to go from raw reads to sequence alignment, genotyping and variant filtering in several simple, reproducible steps.

The pipeline first performs quality trimming on the reads, removing adapter sequences and trimming any bases with a Phred-encoded quality score of less than 30, retaining sequences only with a length greater than 50 bases. Reads are then mapped against the reference genome using bwa mem (Li 2013) and marked for duplicates using Picard (https://broadinstitute.github.io/picard/). The sequences are then realigned using abra2 (Mose *et al*. 2019) to reduce the probability of false positive variant calls around insertions and deletions. Average mapping efficiency was 99.06% (SD: 0.19; min = 98.5, max = 99.44).

The next step in the pipeline performs variant calling at all positions in the genome (i.e. variant and invariant sites) using bcftools (Danecek *et al*. 2021). It parallelises this across 10 Mb windows in the genome to ensure maximum efficiency. Once raw variant calls are complete, the pipeline then performs filtering using vcftools (Danecek *et al*. 2011). Sites are removed if they have calls for less than 80% of individuals, have a Phred quality score of < 30, a minimum depth of 5X coverage and a maximum depth of 15X coverage.

### 2.5. Epigenetic potential quantification

It is important to note that our aim was to identify population-level variation in the presence or absence of CpG sites. Identifying CpG sites in a reference genome sequence is straightforward as they are defined solely by the consensus sequence of bases – in short, they are present or absent in a single reference. However, when considered across a population, CpGs can be present in some individuals but absent in others due to SNPs occurring within CpG sites. Following variant calling, we therefore quantified CpG polymorphism across the genomes of all individuals. To do this, we developed a custom pipeline that operates on each chromosome in parallel. The pipeline first identifies variant positions and then extracts the two flanking sites either side of each position from the reference genome. We then used a custom R script to count the number of CpGs along each chromosome from the flanking sites and variant calls. In brief, this script first identifies all CpGs and also what we call poly-CpGs. CpG variants occur when the reference allele is either a C or G and the flanking sites are G or C respectively. These represent CpG sites that are “lost” with respect to the reference genome in individuals carrying the alternative allele at either the C or G position. Poly-CpG variants occur when the alternative allele is either a C or G and the flanking sites in the reference genome are G or C respectively, and thus represent CpG sites that are “gained” with respect to the reference genome in individuals carrying the alternative allele (as in Hanson *et al*. 2022). The script then identifies the presence or absence of each CpG or poly-CpG site across individuals from which we calculated the total counts per individual. To ensure efficiency, the script works on a per chromosome basis (i.e. it produces chromosome specific totals). Therefore, to calculate genome-wide totals, we used a custom script to combine all chromosome outputs to produce a genome-wide map of CpG positions.

Previous studies calculating epigenetic potential as number of CpGs (e.g. Hanson *et al*. 2020, 2022) have quantified the total number of CpGs present in the specific genome regions and worked at the among-population scale. Therefore, these studies did not account for within-population variation in CpG sites or where in the genome these occur. Because the aim of our study was to identify population-level variation in the presence or absence of CpG sites, we focused on detecting and quantifying the CpG sites that showed variation (i.e. may be present or absent) among our study individuals. However, to rule out the possibility that total CpG counts may yield different results (due to e.g. unaccounted C/G insertions or deletions) we also ran our main models with whole-genome total CpG sites (instead of number of CpG variants). Scripts for epigenetic potential quantification are available at: https://github.com/markravinet/epigenetic_potential

### 2.6. Functional annotation of CpG variants

The GFT-formatted annotation files for the collared flycatcher were converted into BED12 format using the University of California, Santa Cruz (UCSC) gtfToGenePred and genePredToBed programs (available at https://hgdownload.soe.ucsc.edu/downloads.html#utilities_downloads). Variant sites were then annotated using the tool annotateWithGeneParts from the R 4.1.2 package genomation 1.4.1 (Akalin *et al*. 2015). This tool hierarchically classifies the sites into pre-defined functional regions, i.e., promoter, exon, intron, or intergenic. The predefined functional regions were based on the annotation information present in the BED12 files accessed with the genomation tool readTranscriptFeatures. Subsequently, a customized R script was employed to integrate the annotation results of sites with their respective annotation category information.

### 2.7. Quality control analyses

Average sequencing depth was 8.07 (SD: 1.19; min = 5.93, max = 10.52). Due to variation in sequencing depth among variant sites and to increase our confidence in the results, we applied a depth filter to the resulting site variants so that we only included sites with depth ≥ 8x. Note that reported results did not differ from those obtained without applying this filter. Furthermore, we excluded the sex chromosome sequences from our working dataset to avoid biased results due to males having two copies of the Z chromosome as well as due to sex-dependent sequencing quality, as W chromosome was not included in the reference genome and average depth of the Z chromosome highly differed between males and females (mean ± s.d.: 8.50 ± 1.19 vs. 4.86 ± 0.45, respectively), as did average depth between autosomes and Z chromosome, mainly due to the lower depth of female sex chromosome (mean ± s.d.: 8.23 ± 1.18 vs. 6.53 ± 2.02; Fig. S1).

To investigate population structure and general variation among sequenced individuals, we performed principal components analysis on the dataset using PLINK 1.9. We first performed linkage pruning and then PCA on autosomal SNP allele frequencies. Scripts for population structure analyses are available at: https://speciationgenomics.github.io/pca/. Furthermore, because samples were sequenced in two sequencing runs, we used the same PCA to rule out a batch effect (see below).

### 2.8. Statistical analyses

Statistical analyses were carried out in R 4.2.0 (R Core Team 2022), using the “lmer” function of the “lme4” package (Bates *et al*., 2015) to fit mixed models and the “ggplot2” package (Wickham 2016) to develop the figures. All final models fitted assumptions of normality of residuals, normality of random effects, multicollinearity and homogeneity of variance (“check_model” function in “performance” package, Lüdecke *et al*., 2021).

For the two spatial approach comparisons, we ran general linear mixed models with epigenetic potential (CpG + polyCpG count) as the dependent variable. Epigenetic potential was included as dependent variable based on the general prediction that individual categorization according to dispersal propensity would show differences in CpG abundance between group categories, and also allowed us to control by sequencing missingness. Sex, group category (1 vs. 2 or 1 vs. 3, respectively), and their interaction were included as fixed factors, and sampling year was included as a random factor. Percentage of sequencing missingness was included as a weighting variable. The second model (groups 1 vs. 3) was overfitted, and therefore year was removed (see details of sampling year differences below).

For the temporal approach comparison, we ran a general linear model with epigenetic potential (CpG + polyCpG count) as the dependent variable. Sex, group category (1 vs. 4) and the interaction were included as fixed factors. Percentage of sequencing missingness was included as a weighting variable, so that those samples with lower missingness were given more weight. Because all samples from group 4 (first generations of birds in the pine forest) were taken in the same year, the random effect of year was not included in the model. It is important to note that further analyses suggest that neither year nor sequencing missingness drove differences in CpG counts found between the two groups (Fig. S2).

To further test whether the results obtained held depending on CpG type (i.e. CpG vs. polyCpG), CpG location, as well as the patterns followed by other genetic variants, we also ran the same three models described above with i) CpG and polyCpG sites separately, ii) only CpG sites located in promoters or iii) non-CpG variants as dependent variables.

To test whether the observed counts of CpG and polyCpG variants in each annotation category differed from the expected counts, we performed two-sided exact binomial tests using the “stats” package in R. The expected counts were derived by annotating all CpG and poly-CpG sites identified in our analysis.

## 3. RESULTS

### 3.1. Epigenetic potential and its variation in a pied flycatcher population

We identified a total of 801,024 CpG, 735,900 polyCpG and 6,107,566 non-CpG unique variants across the genome. PCA analyses showed no genetic structure associated to dispersal categories (i.e. PCA analysis; Fig. S3), nor sequencing run effect (Fig. S4) within our dataset.

### 3.2. Differences between dispersal group categories

#### Spatial comparison 1: Habitat patch switching

We found a non-significant trend for an effect of the interaction between sex and group category (dispersing or not to a different habitat patch) on epigenetic potential (Group x Sex: F_1,34.23_= 3.685; p= 0.063). Specifically, female dispersers showed a higher number of CpG sites than their non-dispersive counterparts, whereas males showed similar values irrespective of their dispersal status (Males: F_1,15.27_ = 0.006, p= 0.940; Females: F_1,14.69_ = 4.563, p= 0.048; Fig 2, Table 2A).

**Table 2:**
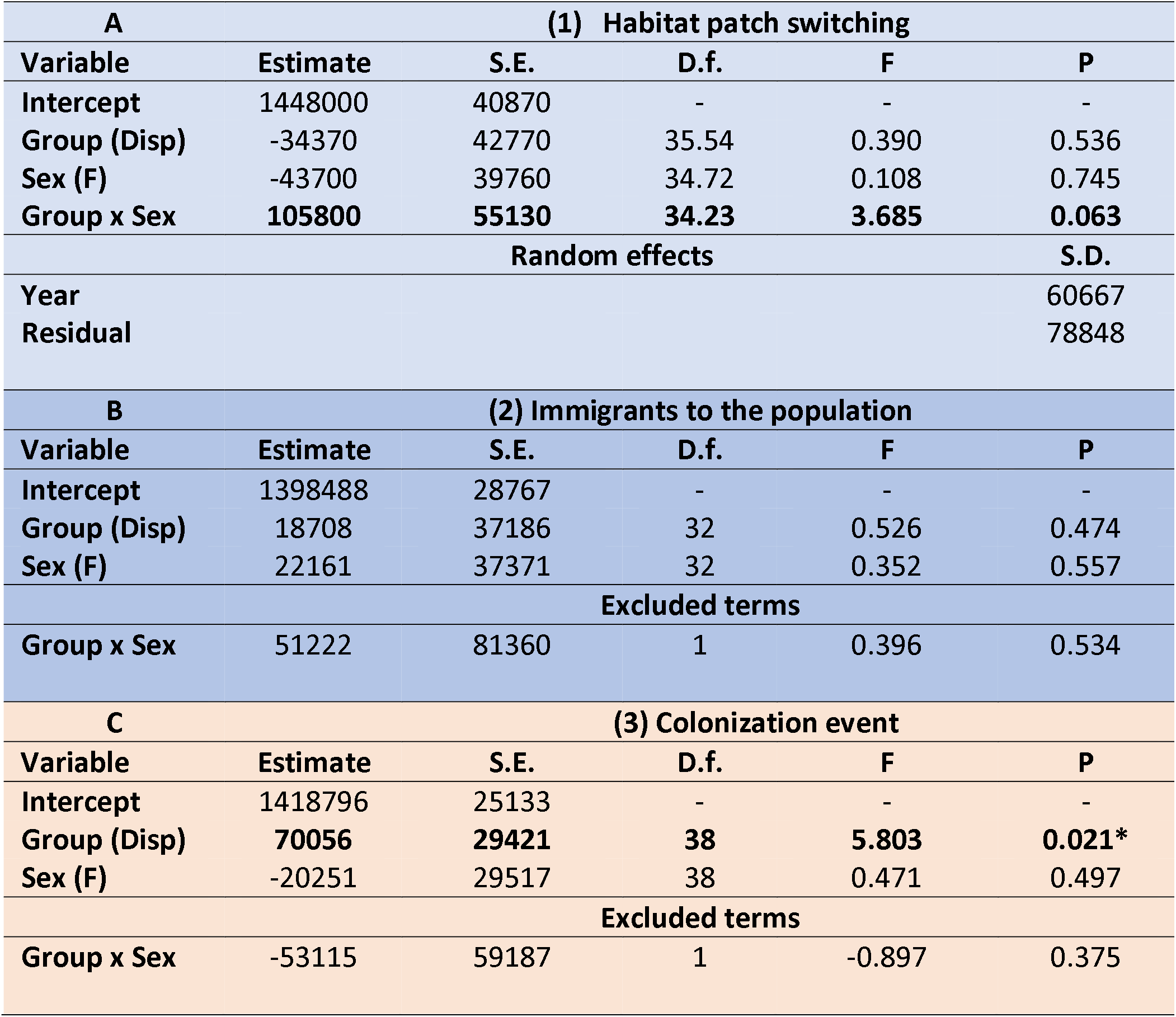
Results of the models testing the association between dispersal group categories and epigenetic potential (quantified as the number of CpG + polyCpG variants across the genome). Table shows the final models corresponding to each of the three comparisons tested, as follows: A (Comparison 1): Habitat patch switching; B (Comparison 2); Immigrants to the population; C (Comparison 3): Colonization event. Note that all three comparisons use the same group of non-dispersers as reference (see Table 1 for details).

**Figure 2.**
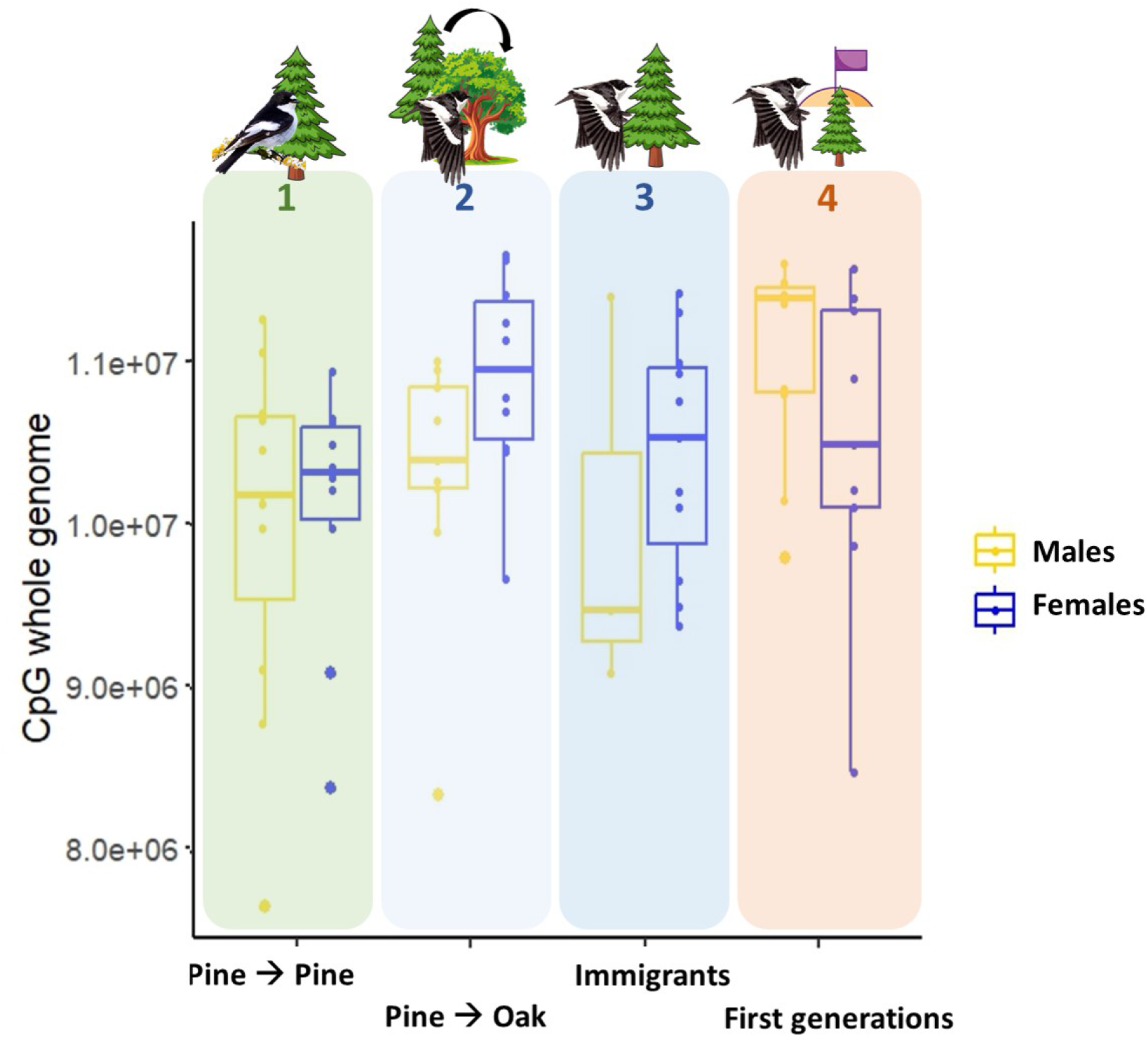
Epigenetic potential (estimated as number of CpG variants across the genome, including CpG and polyCpG sites) across the four group categories included in the study. Group categories correspond to: 1. Birds reared and breeding in the pine forest; 2. Birds reared in the pine forest but breeding in the oak forest; 3. Birds identified as immigrants to the pine population, 4. First generations of birds breeding in the pine forest after patch colonization. The three comparisons tested were made between dispersal categories predicted to have high (blue and red) and low (green) epigenetic potential, and correspond to different spatial (blue vs. green) and temporal (red vs. green) scales. The bottom and top lines of the box represent the interquartile range, and the horizontal line inside the box represents the median. The whiskers represent values outside the lower and upper quartile. Note that CpG counts include total CpG and polyCpG variants pooled, but results were very similar when performing the comparisons for both types separately, and statistical models included sample missingness as weighing variable (see results).

Results did not qualitatively differ when analyzing CpG and polyCpG separately, although the interaction was weaker when considering polyCpGs only (CpG Group x Sex: F_1,34.24_= 3.878; p= 0.057; polyCpG - Group x Sex: F_1,34.27_= 2.703; p= 0.109). Likewise, results did not qualitatively differ when considering CpG sites located in promoters only (Group x Sex: F_1,34.29_= 3.554; p= 0.068), nor when considering non-CpG variants only, although the effect of the interaction was weaker in this case (Group x Sex: F_1,34.34_= 2.546; p= 0.112).

#### Spatial comparison 2: Immigrants to the population

We did not find an association between epigenetic potential and dispersal propensity between individuals categorized as locally-born or immigrants (F_1,32_=0.526, p= 0.474; Fig 2, Table 2B).

Results did not qualitatively differ when analyzing CpG and polyCpG separately (CpG Group: F_1,32_= 0.404; p= 0.530; polyCpG - Group F_1,32_= 1.121; p= 0.298). Likewise, results did not qualitatively differ when considering CpG sites located in promoters only (Group: F_1,32_= 0.001; p= 0.976), nor when considering non-CpG variants only (Group: F_1,22.94_= 1.050; p= 0.315).

#### Temporal comparison: Colonization event

We found a significant association between epigenetic potential and the sampling time within the colonization event (F_1,38_= 5.804; p=0.021). Birds from the first generations after the establishment of the population showed higher number of CpG sites compared to individuals breeding in the same population more than twenty years later (Fig 2, Table 2C).

Results did not qualitatively differ when analyzing CpG and polyCpG separately (CpG Group: F_1,38_= 5.678; p= 0.022; polyCpG - Group F_1,38_= 6.202; p= 0.017), nor when considering non-CpG variants only (Group: F_1,38_= 5.786; p= 0.021). Likewise, results did not qualitatively differ when considering CpG sites located in promoters only, although the association between epigenetic potential and dispersal category was weaker in this case (Group: F_1,38_= 3.995; p= 0.053).

#### Using total CpG count vs. number of CpG variants in the comparisons of epigenetic potential

Results of the main models did not differ overall when using total CpG counts in the whole genome, instead of number of CpG variants (Comparison 1: Group x Sex, p=0.06; Comparison 2: Group, p=0.38; Comparison 3: Group, p=0.008; Fig. S5).

### 3.3. Annotation of CpG variants

As expected, the majority of variants, regardless of type, were located in intergenic regions, followed by introns. Promoters had the fewest CpG and polyCpG variants. This was not the case for non-CpG variants, with the lowest number located in exons (Table S1). Interestingly, distribution of CpG and polyCpG variants across genome regions differed from the distribution of total CpG / polyCpG sites (i.e. including variant and non-variant sites; Table 3; Fig. 3). Specifically, there was a lower percentage of CpG variants located in promoters and exons, and a higher percentage located in introns and intergenic regions, compared to what would be expected if variants were distributed randomly across the genome. In contrast, polyCpGs followed the opposite pattern, with a higher percentage located in promoters and exons, and a lower percentage located in introns and intergenic regions compared to the distribution of total polyCpG sites (Table 3,Fig. 3).

**Table 3:** Results of the binomial test on the differences between the distribution of CpG variants and the distribution of CpG sites (incl. variant and non-variant) across genome regions.

|  | CpG |  | polyCpG |  |
| --- | --- | --- | --- | --- |
|  | p-value | Proportion difference (expected- observed) | p-value | Proportion difference (expected- observed) |
| **Promoter** | < 2.2e-16 | 0.049 | < 2.2e-16 | -0.009 |
| **Exon** | < 2.2e-16 | 0.031 | < 2.2e-16 | -0.011 |
| **Intron** | < 2.2e-16 | -0.060 | 9.682e-05 | 0.002 |
| **Intergenic** | < 2.2e-16 | -0.019 | < 2.2e-16 | 0.017 |

**Figure 3.**
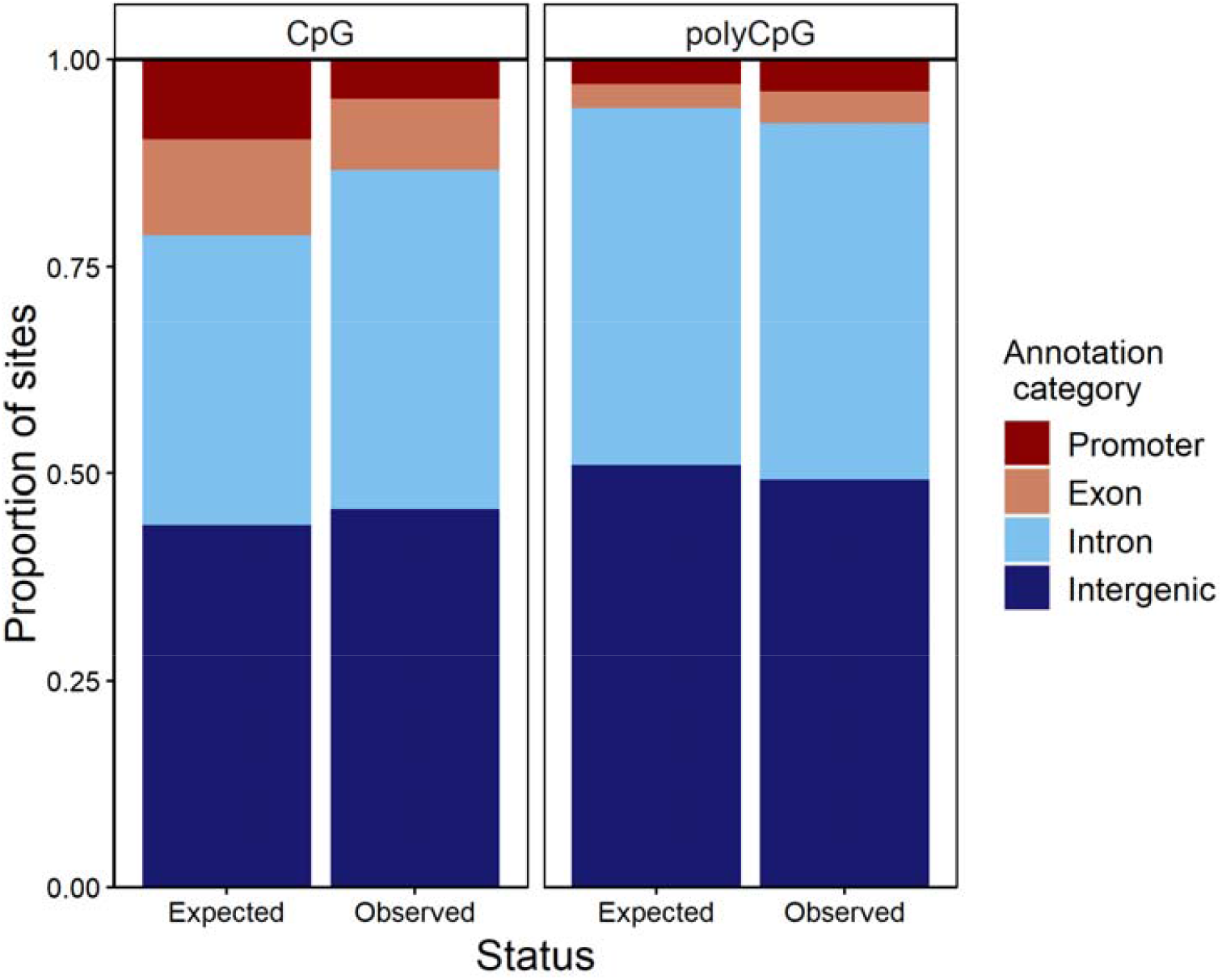
Comparison between the distribution of total CpG sites (incl. variant and non-variant) and CpG variants across genome regions. Expected values correspond to the distribution of total CpG sites across the genome, which would be expected to remain if CpG variants were distributed randomly. Observed values correspond to the actual distribution of CpG variants found in the population (all group categories pooled). For details on the comparisons see methods and Table 3.

## DISCUSSION

Dispersing into a novel environment may require a high degree of phenotypic plasticity that allows individuals to adapt and thrive in the face of environmental changes, and epigenetic mechanisms may be key mediators of these rapid adaptive responses. By performing three comparisons using both spatial and temporal approaches, here we tested whether epigenetic potential for DNA methylation (quantified as the number of CpG variants across the genome) was associated with dispersal behaviour within a bird population. We further functionally annotated CpG variants to test whether their distribution across the genome differed from the distribution of all CpG sites (including variant and non-variant ones).

Epigenetic potential was higher in disperser groups relative to non-disperser groups in two out of the three comparisons. Male and female pied flycatchers from the first generations after the establishment of the population (i.e. first colonizers or their direct descendants; comparison 3; Fig 2), as well as contemporary females dispersing into a non-natal patch to breed (comparison 1; Fig 2) showed higher epigenetic potential compared to individuals from recent generations that remained in their natal patch. This points at an association between CpG variant number and the capacity to move to and/or establish in new environments. This fits our prediction based on the hypothesis that higher number of CpG variants may increase individual susceptibility to show epigenetic changes (i.e., DNA methylation), and thus a plastic response. Furthermore, detecting a sign of this association(s) despite the limitations derived from working on a wild, open population – where controlling for genetic ancestry within our group categories is not feasible beyond origin-based selection criteria (e.g., locally born individuals may have immigrant ancestry) – adds reliability to the pattern. These results set a basis supporting the ecological relevance of epigenetic potential, but a lot remains to be known about the mechanistic links causing this association. For example, whether higher dispersal rates are a direct consequence of higher epigenetic potential, or whether there are other factors underlying this correlation (see below).

We found a strong association between epigenetic potential and dispersal in our temporal approach comparison, with individuals from the first generations after patch colonization showing higher epigenetic potential compared to individuals from recent generations breeding in the same patch. A higher epigenetic potential in expanding vs. stable bird populations has been found previously for an invasive species, the house sparrow (*Passer domesticus*, Hanson *et al*. 2020, 2022), although that study only targeted specific genes of interest. The sparrow work and our findings support a role for epigenetic potential in species establishment success in novel environments. However, because our study does not allow us to infer causality, other factors (e.g., genetic diversity, resilience to stress, different environmental conditions, genetic drift, gene flow) could explain lower numbers of CpG sites in later generations, and therefore further studies and comparisons are necessary to rule these out or determine their relevance.

The association between epigenetic potential and dispersal in pied flycatchers switching habitat patches depended on sex. Females reared in the pine forest but breeding in the oak forest had higher epigenetic potential when compared to site-faithful females. This difference, however, was not observed in males. Previous evidence suggests that the factors driving dispersal and successful settlement may sex-specific (Camacho 2018; Camacho *et al*. 2017b, 2019). For example, in the study population, male dispersal has been previously associated with morphological traits, a pattern that is less evident in females and that disappeared as time passed (Camacho *et al*. 2019). One possibility would be that there is a higher contribution of morphological traits to dispersal in males – due to high competition for breeding territories. This could occur more or less independently on (epigenetically-driven) phenotypic plasticity, which may be more relevant to female dispersal relative to males. Thus, some males, independently on their (epi)genetic traits, may be forced to leave the natal patch due to high levels of competition, whereas only those females with more propensity to settle in novel environments would disperse further. Alternatively, if dispersal itself would entail higher costs for females than for males, females may rely more on (epigenetically-driven) phenotypic plasticity to compensate the costs of dispersal, causing the association between epigenetic potential and dispersal to be stronger or only detectable in females.

Individuals categorized as immigrants to the population did not show differences in epigenetic potential when compared to those categorized as locally-born (comparison 2; Fig. 2; Table 2). One plausible explanation for this result could be the reliability of determining whether an individual is an immigrant (e.g., they may have been born in the surroundings of the study population and/or have ancestors who were born locally in the study area; see above), along with a limited sample size for male immigrants, that could mask the predicted patterns. This possibility could be confirmed or ruled out by genotyping individuals and classifying individuals with genotypes that do not match the majority of the study population as immigrants. Alternatively, immigrants could have ended up in the population by selecting a habitat that resembles the one they were reared in, which would not require phenotypic plasticity (i.e. matching habitat choice; Edelaar *et al*. 2017). This would explain the absence of differences in epigenetic potential between locally-born individuals and those selecting that patch as dispersers. Another plausible explanation would be that immigrants have high epigenetic potential relative to their population of origin, but similar when compared to the values within the population they arrive in. If so, this process could also contribute to the epigenetic potential in the pine forest decreasing over time, via gene flow through short-distance dispersers homogenizing the epigenetic potential values.

The above results overall held when comparing non-CpG variants among dispersal group categories, which could point at the higher number of CpG variants found in some of the disperser groups being a correlate of higher number of total genetic variants. However, we found that CpG variants present in our population were not randomly distributed across the genome, with significant differences compared to what would be expected by chance. Interestingly, these differences followed opposite directions in CpG and polyCpG types, so that CpG variants were more abundant than expected in introns and intergenic regions, and less in promoters and exons. In contrast, polyCpG variants occurred at higher-than-expected frequencies in promoters and exons, and at lower frequencies in introns and intergenic regions. Promoters and exons play crucial roles in gene function, with promoters regulating gene activation and inhibition and exons encoding the transcribed information. As a result, these regions are where most of the genetic variation linked to phenotypic outcomes and adaptation may be expected. Because we used the reference genome of a closely related species (the collared flycatcher) to locate CpG and polyCpG variants, CpG variants (i.e., the ones present in the reference genome, but absent in some of the individuals) could be interpreted as conserved sites from the ancestral lineage. Such conservation might be stochastic or due to sites being potentially essential for normal organismal functioning. In contrast, polyCpGs may better reflect species- or population-specific changes or adaptations to the environment. Some CpG sites might also be highly conserved and therefore CpG variants would be restricted in promoters and exons, as compared to introns and intergenic regions where this variation may not have crucial and / or deleterious phenotypic consequences. PolyCpG variation, on the contrary, may arise and be detected predominantly in regions where such variation leads to phenotypic outcomes and potentially adaptation (i.e. promoters and exons). In mammals, promoter gains and losses are frequent events and have been recognized as key drivers of phenotypic variation (Young *et al*. 2015). If similar mechanisms occur in birds, the observed higher proportion of polyCpG variants in promoters could indeed alter gene expression, promoting phenotypic variation and, subsequently, different levels of phenotypic plasticity. To our best knowledge, this is the first study comparing the distribution of CpG variants vs. totals across different annotation categories throughout the genome. Therefore, results should be interpreted with caution until further research can shed light on the mechanisms underlying these differences or determine whether similar patterns are also found in other species or study systems. Additional research focusing on selection processes acting on specific CpG sites, or comparative studies testing whether population-specific CpG variation is indeed more abundant in regions involved in gene regulation and transcription, are needed to establish the broader ecological relevance of our findings.

Dispersal has been shown to have a genetic component involving many genes (Saastamoinen *et al*. 2018, Hansson *et al*. 2003; Doligez *et al*. 2009; Korsten *et al*. 2013), but also a strong environmental and context dependence (Brown & Brown 1992; Delgado *et al*. 2010). Epigenetic potential is, by definition, a genetic trait and therefore heritable (as it consists of differences in the DNA sequence, i.e. presence or absence of CpG sites). However, the magnitude of the epigenetic variation that will eventually take place within a specific genome will be effectively determined by the environmental context(s) faced by individuals. If epigenetic potential plays a role in the propensity of an individual to disperse, it may be the hub linking genetic and epigenetic inheritance of this complex trait. This would be a key step towards our understanding on the evolutionary impact of both epigenetic potential and dispersal, but also other epigenetic processes. Dispersal decisions scale up into population dynamics and species distributions. Thus, investigating whether the association between epigenetic potential and dispersal also holds at larger spatial (e.g. among populations) and temporal (e.g. species invasions) scales would be a further step towards determining the role of this trait on environmental coping and adaptation.

In light of the above results, a key next step is to investigate the functional links between CpG site density and epigenetically-driven phenotypic plasticity. In this context, the concept of “epigenetic potential” as an individual, genome-wide trait, may to some extent conflict with some of the existing knowledge on how CpG sites and islands affect gene expression. According to previous molecular evidence, not all CpG sites are equally likely to be methylated and / or actively regulate gene expression, with the importance and magnitude of methylation processes in a specific region highly depending on CpG density (Chen *et al*. 2018; Krebs *et al*. 2014; Weber *et al*. 2007), and a high number of closely located CpG sites often sharing methylation status (Weber *et al*. 2007; Bertucci & Parrott 2020). We therefore encourage further research empirically testing test the association between epigenetic potential, (neighboring) CpG density, and variation in gene expression (at CpG site, CpG island, gene and whole genome levels), and eventually whether higher epigenetic potential is associated to (indicators of) phenotypic plasticity. Genetic and biomedical studies have long hypothesized CpG content and density (especially in promoters) to be a key determinant of (epigenetically-driven) variation in gene expression (Yang *et al*. 2014; Edgar *et al*. 2014; Chen *et al*. 2018) with consequences in life history traits, such as lifespan (Sheldon *et al*. 2023; Bertucci & Parrott 2020; Higham *et al*. 2022; McLain & Faulk 2018). Most of this research has been done either within individual genomes or across species, and thus understanding the functional connections and similarities between the previous estimates of CpG density and measures of epigenetic potential (quantified as whole-genome or single-promoter CpG counts) among individuals from the same population remains to be investigated.

Research on epigenetic potential is on its infancy and it remains to be confirmed whether this trait mediates phenotypic plasticity and environmental coping in wild populations. In this context, studies tackling the associations between epigenetic potential and ecologically relevant traits are highly needed to gather empirical evidence that allows us to start delineating and understanding the ecological relevance of this trait. Our results point to an association between epigenetic potential and dispersal within a population at both spatial and temporal scales, becoming the first empirical evidence on the links between genome-wide CpG count and a life history trait in a natural context. This represents a first step towards understanding the relevance of this trait and its potential use as a proxy of individual or population prospects in the face of rapid environmental change.

## DATA AVAILABILITY

Datasets and codes associated to this manuscript will be made available upon acceptance of the manuscript

## AUTHOR CONTRIBUTIONS

Conceptualization: B.J.; Methodology: B.J., J.MP., M.R; Formal analysis: B.J., M.T., M.R; Investigation: B.J., J.MP., D.C., J.P., C.C, J.T.G., J.C.D.; Resources: J.MP., M.R., J.T.G.; Data curation: B.J.; J.MP., M.R., M.T.; Writing - original draft: B.J.; Writing - review & editing: B.J., M.T., M.R., J.MP., D.C., J.P., C.C., J.T.G., J.C.D.; Visualization: B.J., M.T.; Supervision: B.J., M.R., J.MP; Project administration: B.J.; Funding acquisition: B.J., J.MP., M.R., J.T.G.

## ETHICAL NOTE

All applicable international, national and/or institutional guidelines for the capture and ringing of animals were followed and the study was approved by the Spanish institutional authorities. Over the study period, Doñana Biological Station-CSIC and Autonomous Communities of Madrid and Castilla-La Mancha provided capture and ringing licences. Field procedures were approved by the CSIC Ethical Committee for the following projects (refs. CGL2006-07481/BOS, CGL2009-10652, CGL2011-29694, CGL2014-55969-P, PID2019-104835GB-I00, PID2022-141763NA-I00 and PID2022-138133NB-I00) and, more recently, by the Autonomous Community of Madrid (Ref.: PROEX 068.6/24) guaranteeing that they comply with Spanish and European legislation on the protection of animals used for scientific purposes.

## ACKNOWLEDGEMENTS

Many colleagues and friends, too numerous to name individually, helped with data collection since 1988. We are especially grateful to O. Frías, D. Ochoa and F. Romero for their help in the field. We are also grateful for access to the Augusta computing cluster at the University of Nottingham and the Sigma2 Saga computing cluster in Norway. We also thank the Center for Information Technology at the University of Groningen for support and access to the Hábrók high-performance computing cluster.

