## Supplementary Information for "Epigenetic potential and dispersal propensity in a free-living songbird: a spatial and temporal approach"

**Table. S1. Distribution of CpG and not CpG variants across the genome in the study population.**

| Variant status | Annotation category | Counts | Proportion |
|---|---|---|---|
| CpG | promoter | 38484 | 0.048044 |
| | exon | 68109 | 0.085027 |
| | intron | 328158 | 0.409673 |
| | intergenic | 366273 | 0.457256 |
| polyCpG | promoter | 28485 | 0.038708 |
| | exon | 28806 | 0.039144 |
| | intron | 316108 | 0.429553 |
| | intergenic | 362501 | 0.492595 |
| non-CpG | promoter | 214748 | 0.035161 |
| | exon | 164390 | 0.026916 |
| | intron | 2634894 | 0.431415 |
| | intergenic | 3093534 | 0.506508 |

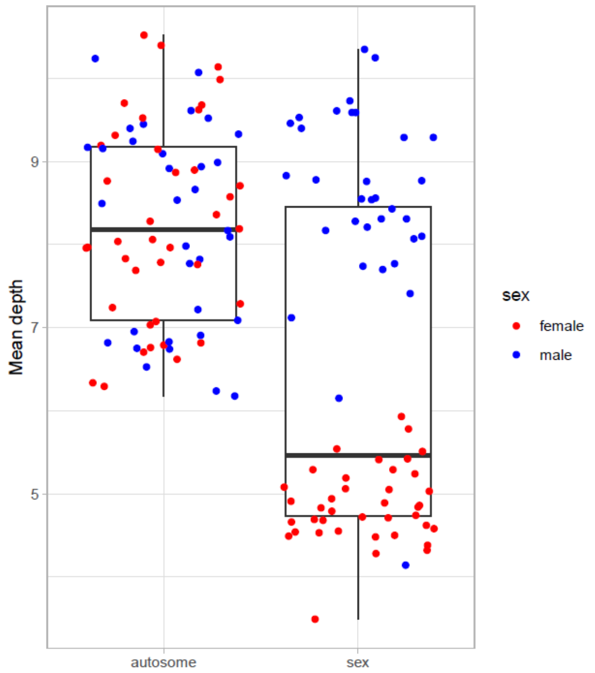


**Fig. S1. Mean sequence depth by sex and chromosome type (i.e. autosomes vs. Z chromosome).**

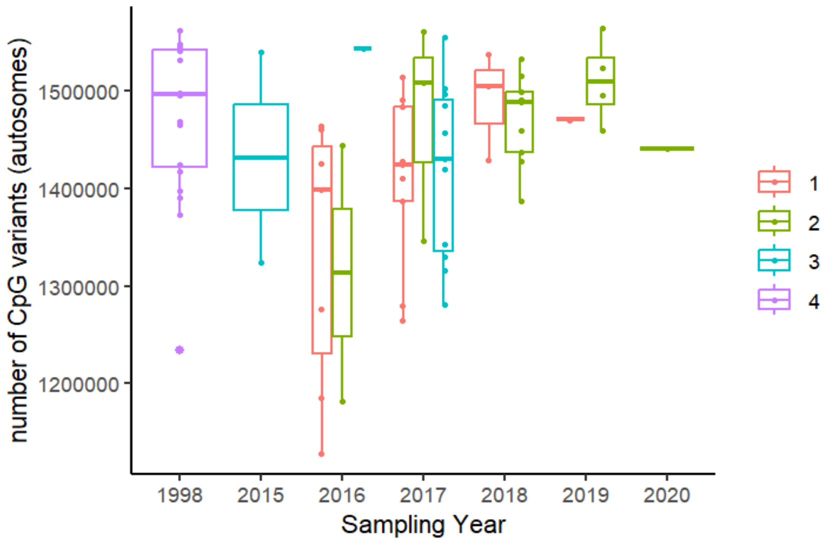


**Fig. S2. Epigenetic potential (calculated as the number of CpG + polyCpG variants) by group category across sampling years. CpG counts in Group 4 sampling year (1998) did not overall differ from CpG counts in those sampling years covered in Group 1. The only significant difference existing among all sampling years was between 1998 and 2016, with lower counts in 2016 (Tukey Test; $p=0.025$). Year 2016 included samples from group categories 1 and 2 and also tended to have lower CpG count compared to other years (i.e. $p < 0.1$ for 2016 vs. 2018 and 2016 vs. 2019) due to higher % missingness. This pattern was confirmed by the above differences in sampling years disappearing when including percentage of sequencing missingness as covariate (Tukey Test; all p values > 0.21).**

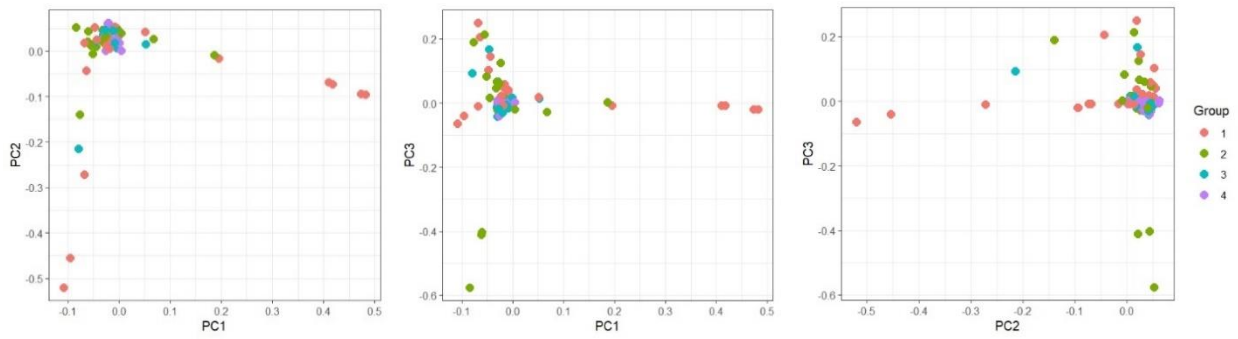


**Fig S3: Principal component analysis showing no genetic variant structure linked to dispersal group categories (for details see methods).**

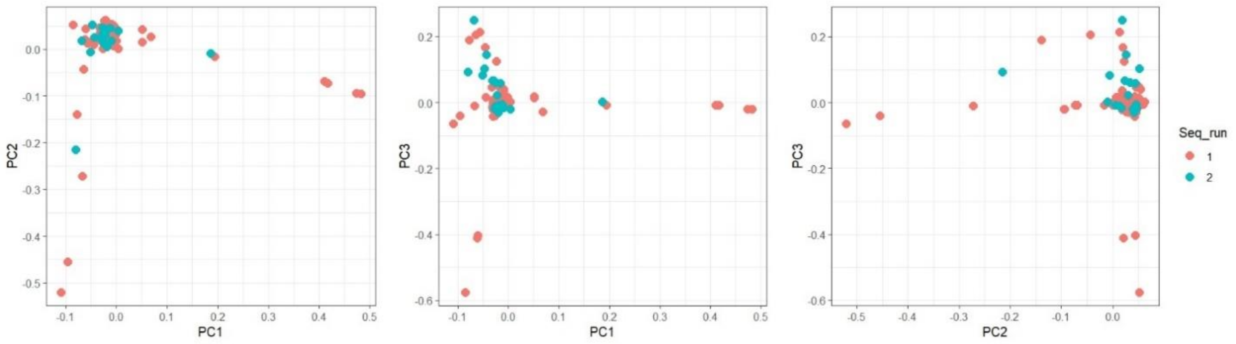


**Fig S4: Principal component analysis showing no genetic variant structure linked to sequencing run (for details see methods).**

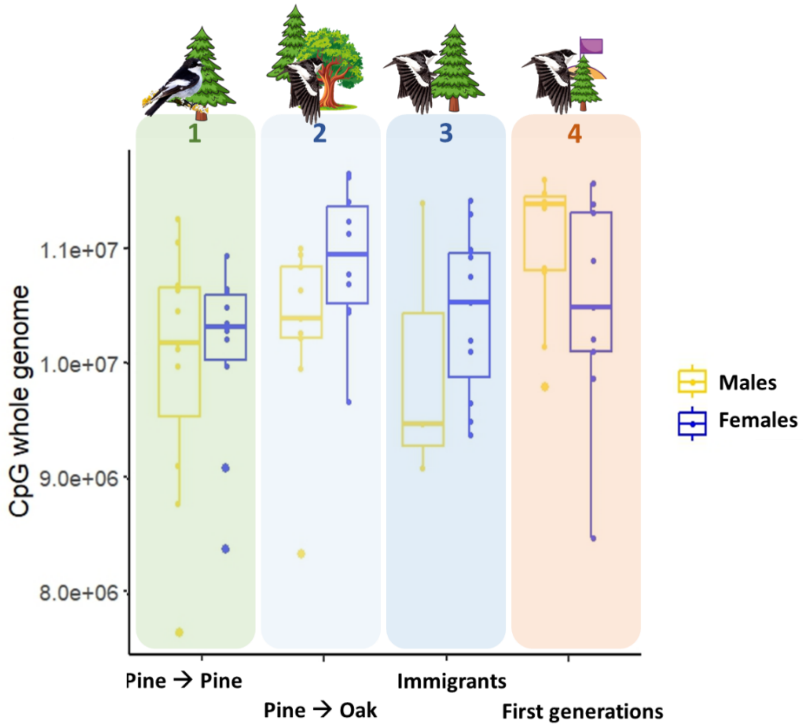


**Fig S5: Epigenetic potential (estimated as total number of CpGs across the genome) across the four dispersal group categories included in the study. Group categories correspond to: 1. Birds reared and breeding in the pine forest; 2. Birds reared in the pine forest but breeding in the oak forest; 3. Birds identified as immigrants to the population, 4. First generations of birds breeding in the pine forest after patch colonization. The three comparisons tested were made between dispersal categories predicted to have high (blue and red) and low (green) epigenetic potential, and correspond to different spatial (blue vs. green) and temporal (red vs. green) scales. The bottom and top lines of the box represent the interquartile range, and the horizontal line inside the box represents the median. The whiskers represent values outside the lower and upper quartile. Note that CpG counts include total CpG and polyCpG variants pooled, but results were very similar when performing the comparisons for both types separately, and statistical models included sample missingness as weighing variable (see results).**
